# Phasic dopamine encodes persistent attraction to reward cues

**DOI:** 10.64898/2025.12.09.693300

**Authors:** Erica S. Townsend, Daniela Garrod, Kyle S. Smith

**Affiliations:** Department of Psychological & Brain Sciences, Dartmouth College, Hanover, NH; Center for Substance Abuse Research, Lewis Katz School of Medicine at Temple University, Philadelphia, PA; Brown University — National Institutes of Health Graduate Partnership Program, Bethesda, MD

## Abstract

Reward-predictive cues can become attractive themselves due to incentive salience attribution. This manifests as persistent engagement with cues, known as sign-tracking. Cue-evoked phasic dopamine in the nucleus accumbens core (NAc) is critical for sign-tracking and cue-evoked reward craving, yet is also known to follow learning rules requiring outcome value updating. Would cue-evoked dopamine track incentive salience persistence, or change in accord with reward value alteration while failing to encode persistent behaviors? We assessed NAc dopamine activity using fiber photometry in sign-tracking rats during an omission task, in which physical cue interaction cancels reward. Animals adapted to reward loss while maintaining non-physical cue interaction reflective of persistent incentive salience. Cue-evoked dopamine remained unchanged during omission, mirroring robust incentive salience but not behavioral adaptations occurring with learning. In contrast, outcome-related dopamine reflected reward gain or loss. Therefore, cue-related dopamine encodes persistence of cue value, independent of outcome-related dopamine signals tracking reward occurrence.

## Introduction

Pavlovian cues that predict rewards help generate motivated behaviors and provide predictive information about outcomes like reward. Often, reward cues can become attractive themselves due to the attribution of incentive salience (Berridge, 2004). This can manifest as physical engagement with stimuli as if it were the reward itself, a response referred to as sign-tracking (Brown & Jenkins, 1968; Flagel et al., 2009; Flagel & Robinson, 2017). Normally, incentive salience and the sign-tracking response closely track changes in reward outcome. Yet, once it is engrained, incentive salience and sign-tracking become divorced from outcome values, such as occurs in cue reactivity in drug addiction (Berridge, 2023; Terry E. Robinson & Berridge, 2025). Thus, it becomes rather impervious to changes in outcome values that would engage updating of the cue value following reinforcement learning rules.

Sign-tracking can come to exhibit habit-like persistence. This is unique as a Pavlovian conditioned response, and is seen in an insensitivity of sign-tracking to stimulus blocking (Holland et al., 2014; María-Ríos et al., 2023), punishment (Chang & Smith, 2016; Townsend et al., 2023), and satiety (Keefer et al., 2022; Kochli et al., 2020). Despite this, sign-tracking can also be highly flexible in some conditions, following model-based learning rules, in which it is sensitive to changes in reward occurrence, homeostatic state, and taste-aversion-based reward devaluation (Amaya et al., 2020; Dayan & Berridge, 2014; Robinson & Berridge, 1993). In general, there appears to be a transition from flexibility in sign-tracking to relative persistence over time.

Dopamine transmission in the nucleus accumbens core (NAc) is required for cue-evoked motivation (Berridge, 2007). This includes a requirement for dopamine in the sign-tracking response (Day et al., 2007; Flagel et al., 2011; Fraser & Janak, 2017; Saunders & Robinson, 2012). Phasic bursts of dopamine release that transition from the time of reward to the time of the cue as Pavlovian cue-reward pairings are experienced (Schultz, 1998) are found selectively in animals that sign-track to the cue rather than pursue the reward during the cue (Flagel et al., 2011). This dopamine activity pattern is therefore consistent with computational models of incentive salience (Flagel et al., 2011; Zhang et al., 2009). Yet, it is also consistent with reinforcement learning computations that confer informational-predictive value to reward cues (Costa & Schoenbaum, 2022; Daw et al., 2005; Dayan & Berridge, 2014; Schultz, 1998). It has remained a point of interest to resolve conditions under which the phasic release of dopamine reflect incentive or predictive value information about reward cues (e.g., Berke, 2018; Berridge, 2007; Garr et al., 2023; Lerner et al., 2021; Mohebi et al., 2019; Smith et al., 2012).

In attempts to decode dopamine roles in motivation versus prediction, motivational signals are typically sought in more temporally long events, with motivation being defined as effort or behavioral activation related to tonic or ramping dopamine levels (Enriquez-Traba et al., 2024; Hamid et al., 2016; Howe et al., 2013; Mohebi et al., 2019; Niv et al., 2007; Salamone et al., 2007). However, motivation can separately be generated by phasic spikes of activity to reward cues in limbic systems, which take motivation levels from low to high immediately by virtue of incentive salience (Ahrens et al., 2016; Berridge, 2004; Day et al., 2006; Ferguson et al., 2020; Flagel et al., 2011; Fraser et al., 2025; Smith et al., 2011; Wyvell & Berridge, 2000). There remains a challenge in understanding phasic dopamine function for learning and motivation. This challenge is particularly relevant for understanding what role phasic dopamine plays when cue-triggered motivations persist in a habit-like manner despite decrements in the value of the outcome that these cues predict. This is thought to be a component of addiction, in which the power of reward cues to elicit craving and motivate behavior persists and becomes decoupled from the actual values of the outcome (Edwards & Koob, 2010; Robinson & Berridge, 2008; Saunders & Robinson, 2013; Tunstall & Kearns, 2015).

To gain entryway to this issue, we measured dopamine activity dynamics as animals acquired sign-tracking behavior to a reward-predictive lever cue, and then encountered an omission contingency. During omission, sign-tracking behavior that results in physical cue engagement (deflection of a lever that signals reward as a Pavlovian cue) is punished by reward loss. Animals are slow to reduce their responding to avoid reward cancellation, and never fully stop (Breland & Breland, 1961; Chang & Smith, 2016; Davey et al., 1981; Locurto et al., 1976; Stiers & Silberberg, 1974; Townsend et al., 2023; Townsend & Smith, 2025; Williams & Williams, 1969). Notably, even when animals do adapt to the new contingency, they still do not stop sign-tracking; rather, they change their repertoire of behavior by reducing movements that lead to lever deflection and increasing movements that do not (Chang & Smith, 2016; Locurto et al., 1976; Townsend et al., 2023; Townsend & Smith, 2025). Therefore, the omission contingency procedure produces a favorable situation for assessing dopamine function in persistent incentive salience and learning: animals are learning from reward loss and adapting their behavior accordingly, yet they are maintaining a robust attraction to the cue.

## Results

### Omission procedures do not alter the incentive value of the cue

We first used two groups of animals to confirm that the reward-predictive cues maintain their incentive value across omission learning days. All animals were trained over 12 sign-tracking acquisition sessions to associate the insertion of a CS+ lever cue with subsequent reward delivery. One group was then given a conditioned reinforcement (CR) assay in which a novel lever was presented, pressing of which caused the presentation of the CS+ (Figure 1a). A second group was trained and then exposed to an omission contingency over 5 additional sessions, followed by CR testing. During omission, any physical engagement with the CS+ lever that led to its deflection resulted in reward cancellation. Both groups initially acquired sign-tracking behavior similarly, reflected in frequent engagement with the CS+ lever that resulted in its deflection (Figure 1b). The group that was then exposed to the omission contingency reduced sign-tracking behaviors that led to lever deflection (Figure 1c). As seen previously (Chang & Smith, 2016; Townsend et al., 2023; Townsend & Smith, 2025), there was a reduction in CS+ deflections that occurred gradually over 5 days (125 trials). CS+ deflections never fell to zero, reflective of the CS+ carrying a markedly strong motivational pull. Nevertheless, animals did learn to mostly avoid lever deflections due to reward cancellation. The reduction in lever deflections over days was equally driven by learning from rewarded and unrewarded trials, as pressing during trials following reward receipt vs. reward cancellation were similar (see Supplemental Figure 1). In CR testing, the reward-paired CS+ was a similarly effective reinforcer for both groups of animals: both groups had a high number of presses on the CR lever resulting in many CS+ presentations (Figures 1d-f). Sign-tracking to the CS+ when it was earned was also similar between the animal groups (Figure 1f-g). Thus, omission exposure did not reduce the incentive value of the cue at all.

**Figure 1:**
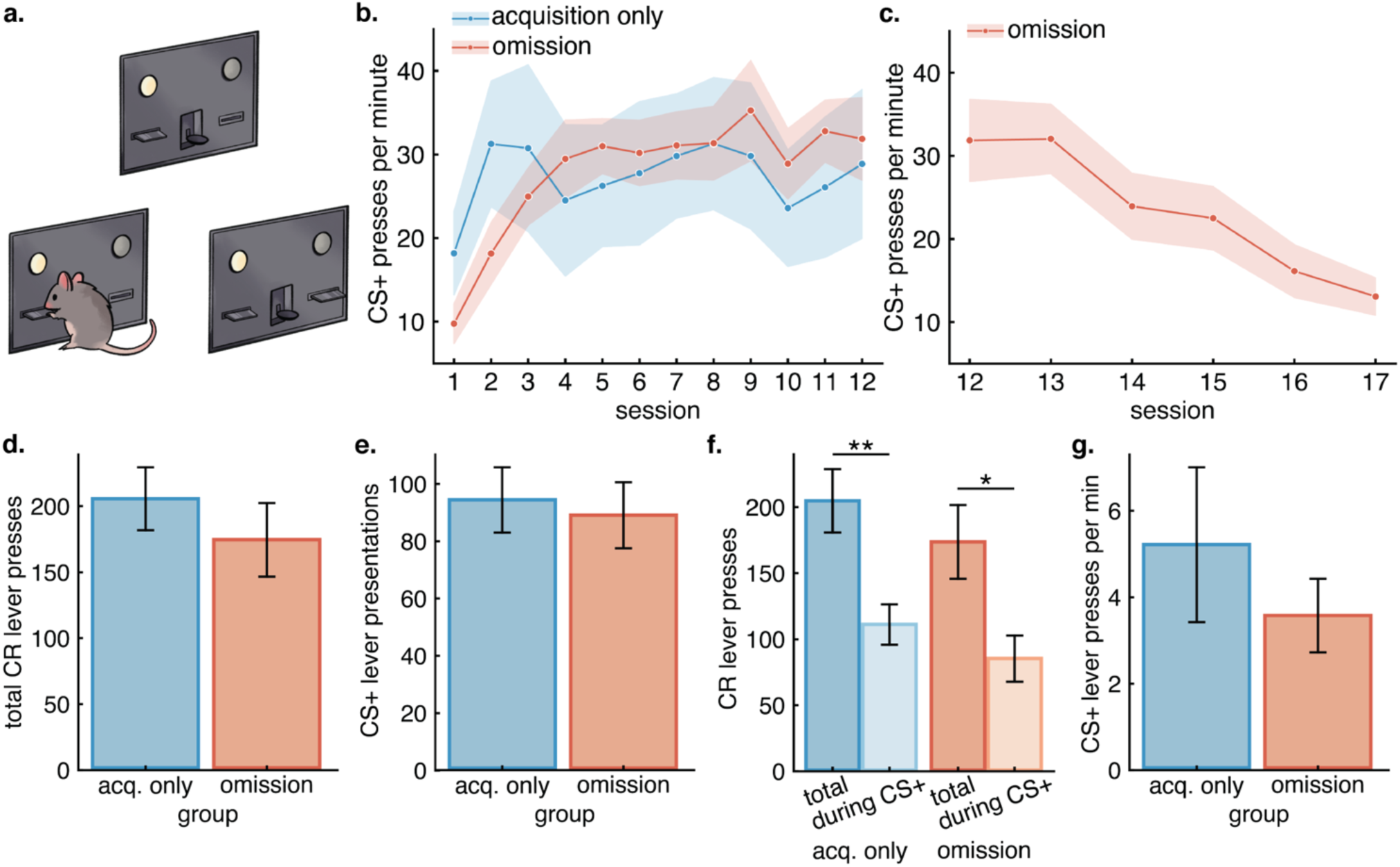
Conditioned reinforcement. **A)** Schematic of the conditioned reinforcement task. **B)** CS+ lever presses per minute across the 12 sign-tracking acquisition training sessions for animals in the acquisition only (blue) and omission (scarlet) groups. Press rates between groups were similar on the final session (t=-0.294, p=0.773) despite acquiring sign-tracking at different rates (no effect of session or group; significant interaction between session and group: est: 1.29, CI: 0.46 – 2.12, p=0.003). **C)** CS+ lever presses per minute for animals in the omission condition across all 5 omission sessions. Pressing significantly decreased over sessions (est: -4.09, CI: -5.35 – -2.82, p<0.001). **D)** Total CR lever presses during the testing session for animals in the acquisition only (blue) and omission (scarlet) groups. **E)** Number of CS lever presentations during the testing session for animals in the acquisition only (blue) and omission (scarlet) groups. **F)** CR lever presses over the total session (darker shades) and only during CS+ presentations (lighter shades) for acquisition only animals (blue; t=3.29, p=0.005) and omission animals (scarlet; t=2.68, p=0.018). **G)** CS+ lever press rates during the testing session for animals in the acquisition only (blue) and omission (scarlet) groups. For all plots, error bars and ribbons indicate ±SEM, and asterisks (*) represent statistical significance.

### Dopamine recording during sign-tracking acquisition and omission: behavioral characterization

A separate set of animals were given intra-NAc viral injections containing the D2-receptor based green fluorescent GRAB_DA2m_ indicator (Sun et al., 2020) and implanted with optic fibers for photometry recording. The animals underwent 12 sessions of sign-tracking acquisition training followed by omission (Figure 2a). In this experiment, there was a second CS- lever interspersed with the CS+ lever, which predicted nothing; engagement with the CS- was always minimal (Supplemental Figure 2). Sign-tracking to the CS+ was acquired over the 12 sessions as in the conditioned reinforcement experiment. During these sessions, readouts of CS+ lever deflections increased and became faster to occur, while food-cup entries during the CS+ decreased (Figure 2b-e, teal).

**Figure 2:**
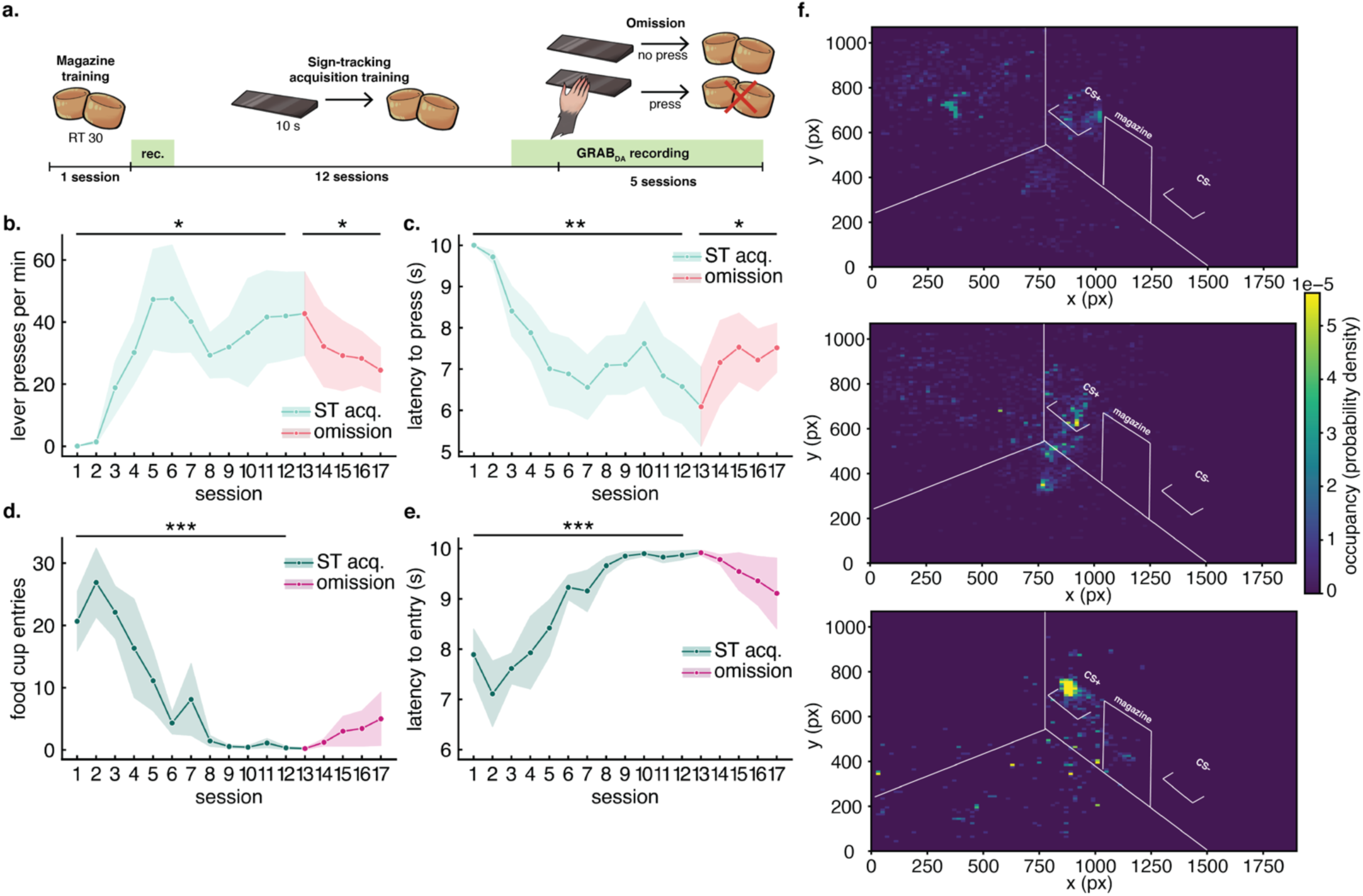
Lever-directed behaviors during sign-tracking acquisition and omission. **A)** Schematic of task. **B)** CS+ lever presses per minute during sign-tracking acquisition training (sessions 1-12; teal) increased as animals acquired the response (effect of session: est: 0.75, CI: 0.16 – 1.34, p=0.013) and decreased over omission (sessions 13-17, pink) as the contingency was learned (effect of session: est: -0.92, CI: -1.78 – -0.06, p=0.037). **C)** Latency to press in seconds during sign-tracking acquisition training (sessions 1-12; teal) decreased across sessions (effect of session: est: -0.26, CI: -0.41 – -0.10, p=0.001) and increased during omission (sessions 13-17, pink; effect of session: est: 0.24, CI: 0.03 – 0.45, p=0.024). **D)** Food cup entries during CS+ presentations throughout sign-tracking acquisition training (sessions 1-12; deep teal) decreased significantly over sessions (effect of session: est: -2.26, CI: -2.96 – - 1.57, p<0.001), remaining low through omission (sessions 13-17, magenta). **E)** Latency to food cup entries during CS+ presentations through sign-tracking acquisition training (sessions 1-12; teal) increased significantly over sessions (effect of session: est: 0.24, CI: 0.17 – 0.32, p<0.001). During omission, food cup entry latency remained relatively unchanged. **F)** Representative location heatmaps during CS+ lever presentations for sign-tracking acquisition session 1 (top), 12 (middle) and average over omission session (bottom). For all plots, error ribbons represent ± SEM. Asterisks (*) represent significant results.

Sign-tracking involves a complex behavioral repertoire of CS+ orienting, sniffing, grabbing, and biting. We have previously shown that omission exposure leads to a restructuring of this behavioral repertoire but that sign-tracking overall is maintained in subtler ways that do not result in physical CS+ engagement and reward loss (Chang & Smith, 2016; Townsend et al., 2023; Townsend & Smith, 2025). In contrast, we have found that extinction exposure leads to a loss of sign-tracking entirely, despite the reduction in lever deflections looking identical between extinction and omission conditions (Townsend & Smith, 2025). Further, this behavioral pattern does not occur in animals whose reward experience is yoked to animals undergoing omission learning (Chang & Smith, 2016), suggesting that it is not simply a reflection of changes in reward delivery likelihood.

This omission contingency, identical to the above experiment on conditioned reinforcement, was imposed on the animals for 5 days after the end of sign-racking acquisition. As seen in the above experiment, and in the literature (Chang & Smith, 2016; Townsend et al., 2023; Townsend & Smith, 2025), CS+ deflections were slow to reduce and never fully ceased, despite many experiences with reward loss. Reward losses on each of the 5 omission days were, on average, 15, 14, 10, 12, and 11 trials in which animals could not withhold their physical CS+ interaction and reward was not provided as a consequence. Still, animals did learn over time to adjust their behavior.

There was a decrease in CS+ lever deflections and an increase in the latency to deflect the CS+ (Figures 2b-c, pink), reflective of learning from reward loss. Food cup entries during the CS+ remained quite low; however a modest and nonsignificant trend of increasing food cup engagement was observed, akin to a “checking” behavior previously reported in a similar omission paradigm (see Townsend et al., 2023; Figures 2d-e, pink).

Video analysis of behavior and head position with respect to the CS+ can capture sign-tracking presence or absence (Iglesias et al., 2023; Townsend & Smith, 2025).

Therefore, we performed video analyses using DeepLabCut to visualize the positioning of animals’ heads in the chamber during CS+ presentations and found that CS+ attraction was considerable in all animals. This analysis indicated similar head positioning during 10-seond CS+ presentations during the final acquisition session and throughout omission, suggesting animals did not cease their lever attraction despite the decline in pressing (Figure 2f).

### DA release dynamics change in accord with sign-tracking acquisition

There was a clear and expected evolution in dopamine release patterns from the first to the final sign-tracking acquisition sessions (Figures 3c-d). CS+ evoked dopamine release spikes grew by a marked 784% from session 1 to 12. Dopamine release to the reward pellet feeder click and to the reward delivery remained relatively lower and unchanged. This was evident when either the area-under-the-curve (AUC) or peak z-scores were considered of CS+ dopamine release events (Figures 3e-h). These findings, which are consistent with prior results (Day et al., 2007; Flagel et al., 2011), show that the development of sign-tracking to a reward cue coincides with an accentuation of phasic dopamine activity selectively at the onset of that cue. There was not a correlation between levels of dopamine elicited by the CS+ and the number of CS+ deflections (Figure 3i), thus it is unlikely that transmission at cue onset is due to vigor and applied force. Similarly, dopamine release was strongly aligned to CS+ lever insertion that triggered sign-tracking, and not to the lever deflections resulting from sign tracking (Figure 3j).

**Figure 3:**
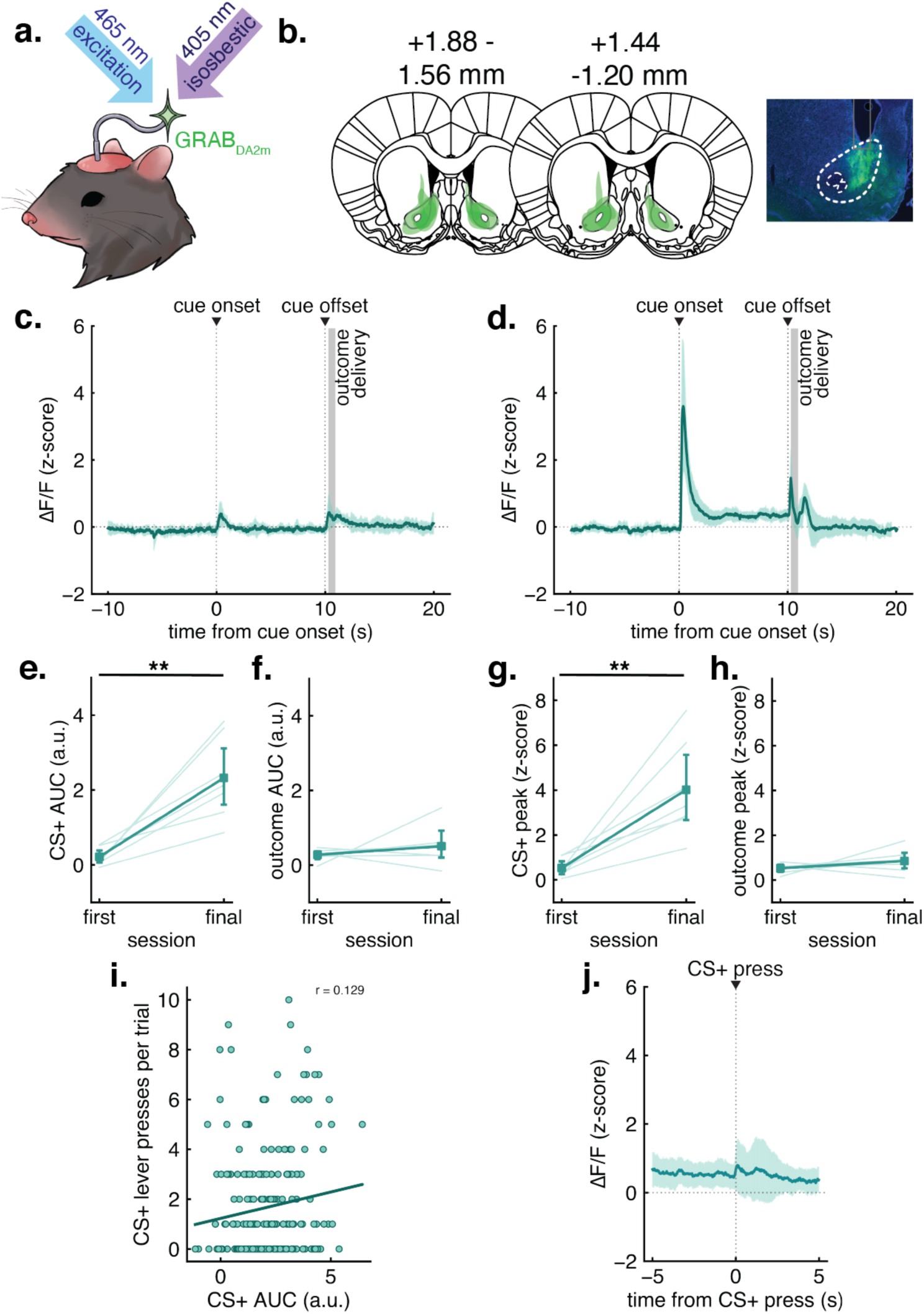
DA responses during sign-tracking acquisition. **A)** Schematic of *in vivo* fiber photometry measurement. **B)** Viral mapping of GRAB_DA_ (GFP) in the NAc. **C)** Average ΔF/F of the 25 CS+ trials in the first sign-tracking acquisition session for the 10 s prior, 10 s during, and 10 s following the cue presentation. Cue onset (0 s) and cue offset (10 s) are noted by dotted lines. Outcome delivery is noted by a shaded gray area beginning at the moment of the first pellet’s release from the hopper, ending when the second is released. **D)** Average ΔF/F of the 25 CS+ trials in the final sign-tracking acquisition session for the 10 s prior, 10 s during, and 10 s following the cue presentation. **E)** AUC (arbitrary units; a.u.) of DA elicited by the cue was greater on the final session (t=4.887, p=0.003). **F)** AUC of DA elicited by pellet delivery following the cue. **G)** Peak z-score during the 1 s time bin beginning at the cue onset was greater on the final session (t=4.242, p=0.005). **H)** Peak z-score after outcome delivery, in a 1 s time bin beginning 1 s after cue offset. **I)** No correlation between AUC calculated during the 1 s cue onset bin and the number of lever presses during given trials in the final session of sign-tracking acquisition for each animal. For all plots, error ribbons and bars represent ± SEM. Asterisks (*) represent significant results.

Dopamine signaling at the CS- was stably low across sessions. CS- evoked dopamine exhibited a modest increase from baseline, but it quickly and sharply decreased to below baseline following CS- lever insertion and retraction (Supplemental Figure 3). Peak dopamine responses evoked by the CS- was only 26.4% of dopamine evoked by the CS+ by the final session of training; this relatively low CS- signal tracked the low and unchanged behavioral interaction with the CS- (Supplemental Figures 2-3).

### Divergent DA dynamics reflect omission learning and motivational persistence

Dopamine recordings then continued daily as animals were exposed to, and learned, the new omission contingency (Figure 4a). Dopamine release at the time of CS+ delivery during omission learning remained stable (Figure 4b-d). The AUC and peak z-score of dopamine elicited by the CS+ remained robust and stable throughout the omission period. CS+-evoked dopamine was not distinguishable between cancelled (Figure 4e, g) or rewarded trials (Figure 4f, h). Thus, the stable dopamine levels at the CS+ onset tracked the stable sign-tracking behavior and stable incentive value of the cue (as noted in the conditioned reinforcement experiment as well). Cue-evoked dopamine did not change whatsoever based on learning from reward occurrence or cancellation as might be expected the phasic CS+ dopamine signal reflected reward prediction.

**Figure 4:**
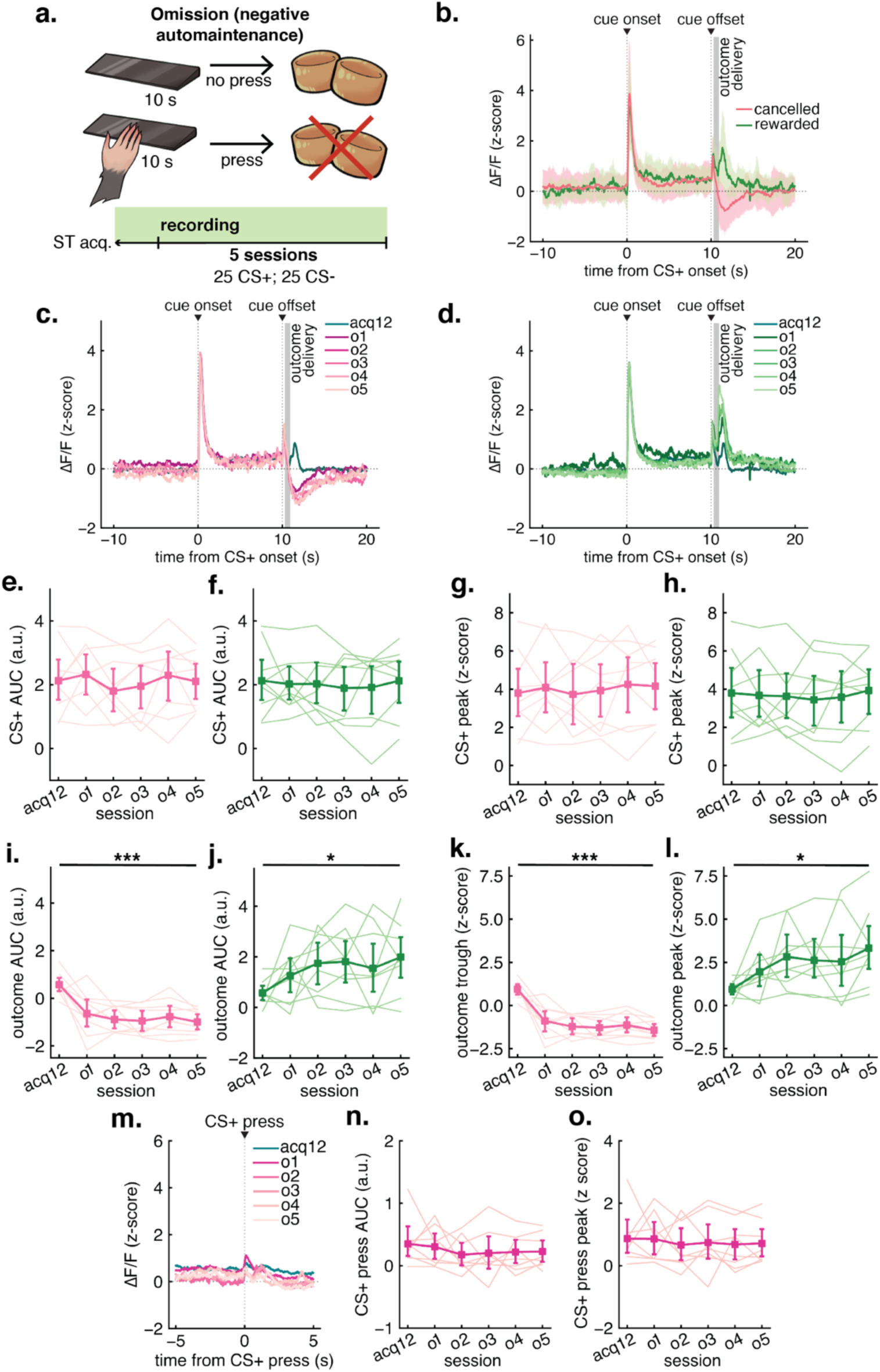
DA responses during omission. **A)** Schematic of the omission schedule. **B)** Average ΔF/F of the cancelled (pink) and rewarded (green) CS+ trials in the first omission session for the 10 s prior, 10 s during, and 10 s following the cue presentation. Cue onset (0 s) and cue offset (10 s) are noted by dotted lines. Outcome delivery is noted by a shaded gray area beginning at the first pellet’s release from the hopper, ending when the second is released. **C)** Average ΔF/F of the 25 CS+ trials in the final sign-tracking acquisition session (teal) and cancelled trials in each omission session (pink, darkest to lightest in color from sessions 1-5) for the 10 s prior, 10 s during, and 10 s following the cue presentation. **D)** Average ΔF/F of the 25 CS+ trials in the final sign-tracking acquisition session (teal) and rewarded trials in each omission session (green, darkest to lightest in color from sessions 1-5) for the 10 s prior, 10 s during, and 10 s following the cue presentation. **E)** AUC (arbitrary units; a.u.) of DA elicited by the cue in the final sign-tracking acquisition session (acq12) and cancelled trials in each subsequent omission session (o1-o5). **F)** AUC of DA elicited by the cue in the final sign-tracking acquisition session (acq12) and rewarded trials in each subsequent omission session (o1-o5). **G)** Peak z-score during the 1 s time bin beginning at the cue onset in the final sign-tracking acquisition session (acq12) and cancelled trials in each subsequent omission session (o1-o5) decreased over sessions (effect of session: est: -0.36, CI: -0.50 — -0.23, p<0.001). **H)** Peak z-score during the cue in the final sign-tracking acquisition session (acq12) and rewarded trials in each subsequent omission session (o1-o5) increased over sessions (effect of session: est: 0.39, CI: 0.10 – 0.68, p=0.010). **I)** AUC of DA elicited by the outcome in the final sign-tracking acquisition session (acq12) and cancelled trials in each subsequent omission session (o1-o5). AUC decreased over sessions (effect of session: est: -0.24, CI: -0.34 – -0.14, p<0.001). **J)** AUC of DA elicited by the outcome in the final sign-tracking acquisition session (acq12) and rewarded trials in each subsequent omission session (o1-o5). AUC increased as the task progressed (effect of session: est: 0.23, CI: 0.05 – 0.41, p=0.013). **K)** Trough z-score after outcome delivery, in a 1 s time bin beginning 1 s after cue offset, in the final sign-tracking acquisition session (acq12) and cancelled trials in each subsequent omission session (o1-o5). **L)** Peak of DA elicited by the outcome in the final sign-tracking acquisition session (acq12) and rewarded trials in each subsequent omission session (o1-o5). **M)** Average ΔF/F of all CS+ lever presses in the final sign-tracking acquisition session (teal) and each omission session (pink, darkest to lightest in color from sessions 1-5). **N)** AUC of DA transmission 0.5 seconds from CS+ lever presses during trials in the final sign-tracking acquisition session (acq12) and in each subsequent omission session (o1-o5). **M)** Peak DA transmission 0.5 seconds from CS+ lever presses during trials in the final sign-tracking acquisition session (acq12) and in each subsequent omission session (o1-o5). For all plots, error ribbons and bars represent ± SEM. Asterisks (*) represent significant results.

Immediately in the first omission session, a pronounced distinction appeared in outcome-related dopamine dynamics between cancelled and rewarded trials. Dopamine activity decreased below baseline when no pellets were delivered due to CS+ deflections. On rewarded trials, in which the CS+ was not deflected, outcome signaling remained above baseline and higher than signal that occurred on the last acquisition session (Figure 4b). This pattern continued across the entirety of the 5-day omission learning period (Figure 4c-d), in which reward loss reliably coincided with dopamine dips (Figure 4i, k) and reward gain coincided with dopamine rises (Figure 4j-l). Despite these alterations in dopamine activity at the outcome relating to whether animals deflected the CS+ lever or not, dopamine did not correspond to CS+ lever pressing (Figure 4m), nor did press-related signaling change in magnitude across sessions (Figure 4n-o).

### Event-related dopamine alterations do not occur rapidly trial-to-trial

As outcome-evoked dopamine dipped to reward loss and rose to reward gain at the learned time of reward delivery, we next examined if the intensity of event-related signaling events were related to whether rewards had occurred or not on the previous trial. They were not. CS+ evoked dopamine remained robust without relation to current or previous trial history during both cancelled (Figures 5a, c) and rewarded trials (Figures 5b, d). This is in line with stable CS+ transmission seen across sessions.

**Figure 5:**
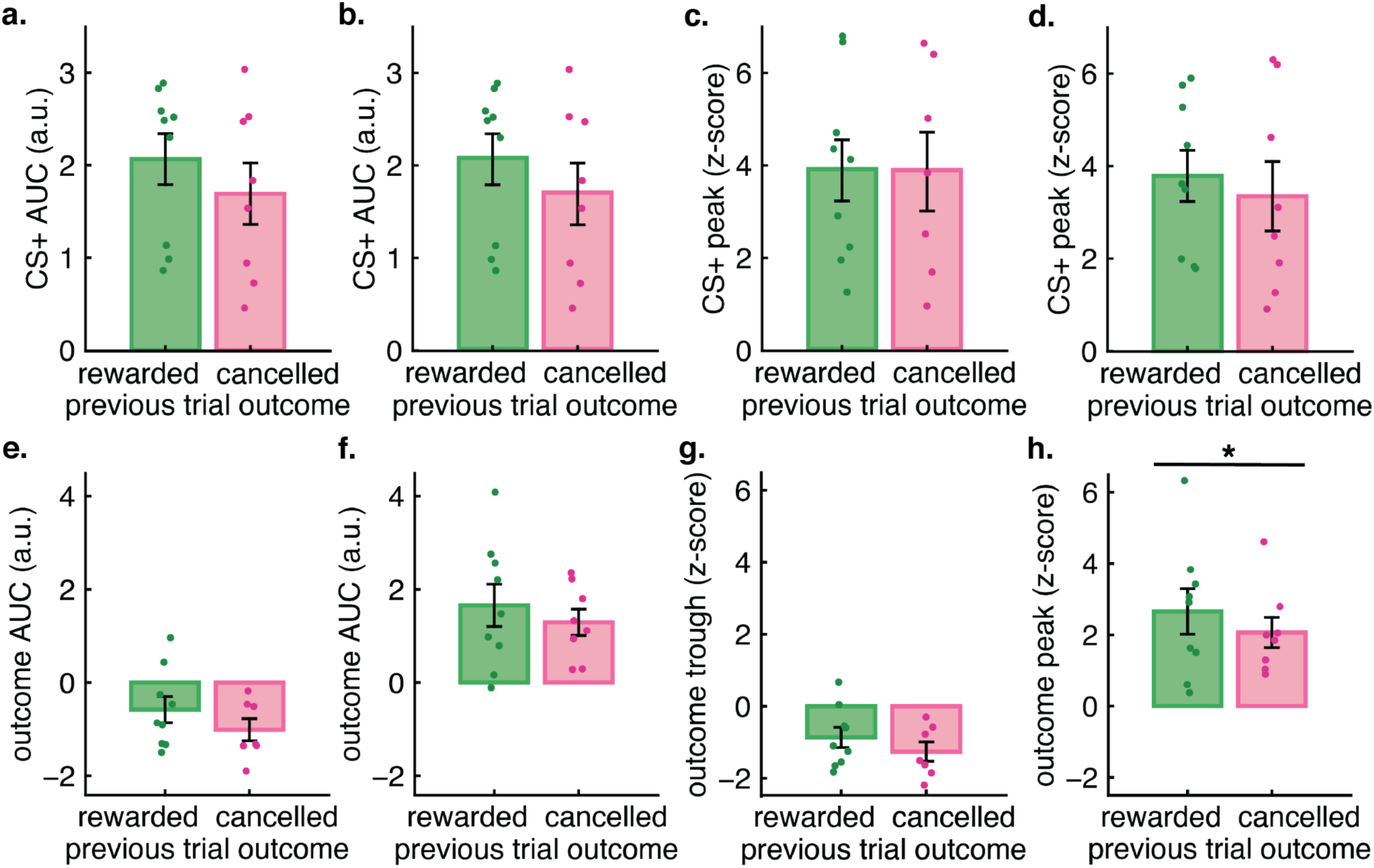
Trial-by-trial analysis of CS+ and outcome evoked DA signaling. **A)** AUC of CS+ responses during cancelled trials when the previous trials were either rewarded or cancelled. **B)** AUC of CS+ responses during rewarded trials when the previous trials were either rewarded or cancelled. **C)** Peak z-score of CS+ responses during cancelled trials when the previous trials were either rewarded or cancelled. **D)** Peak z-score of CS+ responses during rewarded trials when the previous trials were either rewarded or cancelled. **E)** AUC of outcome responses following cancelled trials when the previous trials were either rewarded or cancelled. **F)** AUC of outcome responses following rewarded trials when the previous trials were either rewarded or cancelled. **G)** Trough z-score of outcome responses following cancelled trials when the previous trials were either rewarded or cancelled. **H)** Peak z-score of outcome responses following rewarded trials when the previous trials were either rewarded or cancelled; this was the only significant difference (t=2.64; p=0.034), but was moderate and not present in AUC analysis. For all plots, error bars represent ± SEM. Asterisks (*) represent significant results.

Intriguingly, despite cross-session dynamics in dopamine release at the outcome time (reward or loss of reward), the strength of outcome-related dopamine dips on cancelled trials were unrelated to reward occurrence following the previous trial (Figure 5e, g), and gradual session-wide rises of outcome-evoked dopamine on rewarded trials were also generally unrelated to what occurred on the last trial (Figure 5f, h). Therefore, changes in dopamine representation of the outcome reflected the session-wide experience rather than of a potential trial-by-trial comparison.

## Discussion

Collectively, these results support a role for phasic dopamine activity in persistent, yet flexible, motivational reactions to reward cues. Phasic dopamine signaling tracked the robust behavioral attraction to the cue and did not change as rewards and reward losses occurred, even as such reward losses – a considerable 125 omission trial exposures with 62 average reward losses – led to behavioral learning.

The cue-evoked dopamine was fully independent of changes in dopamine release that occurred during the outcome receipts or losses themselves. Outcome-related dopamine activity seen here could reflect a process of “updating” the incentive value of rewards and their paired cues (McClure et al., 2003; Robinson & Berridge, 2025; Zhang et al., 2009). Such updating would normally translate to corresponding changes in the incentive qualities of the cue, which is consistent with dopamine changes observed during sign-tracking acquisition (see Figure 3).

In our omission test, however, such updating did not affect incentive properties of the cue, nor did it affect cue-evoked dopamine. The omission conditions therefore engendered a persistent motivational attraction to cues in a manner that could mimic what is proposed to occur in addiction, in which the motivational draw of cues becomes decoupled from such updating (Robinson & Berridge, 2025). Ultimately, this leads to cue attraction that is persistent in the sense of being in relative disregard to remembered or received outcomes. “Irrational” motivation of this sort can be extremely powerful in guiding behavior, such as occurring even when outcome is remembered as fully aversive (Berridge, 2023; Robinson & Berridge, 2013; Warlow et al., 2020).

One could instead conclude from these results that cue-related dopamine release reflects incentive salience, while reward-related dopamine separately reflects prediction errors. Several studies report distinctions in motivation and reinforcement learning through differing dopamine dynamics (Enriquez-Traba et al., 2024; Goedhoop et al., 2023; Hamid et al., 2016; Mohebi et al., 2019). Motivationally relevant signals are often pinned to be temporally drawn-out tonic or ramping dopamine events. However, we observed no dopamine ramping (Hamid et al., 2016; Howe et al., 2013; Mohebi et al., 2019), no detectable tonic changes (Mohebi et al., 2019), nor a relationship of dopamine with sign-tracking vigor (i.e., lever deflection) that might reflect an effort characteristic of motivation that is known to involve accumbens dopamine (Salamone et al., 2007). Still, the possibility of separable cue-related incentive salience and outcome-related reward prediction error encoding deserves to remain on the table.

While dopamine responses to outcomes under omission conditions initially appeared as an error-like signal, this was less clear at the cross-session level. Perhaps when observing only within smaller temporal windows, such as a given hour long session, these signals appear more as a standard prediction error. When multiple sessions are considered though, the intensification of error-like signals may reveal an additional underlying mechanism, such as growing uncertainty surrounding reward receipt. Uncertainty in reward occurrence can encourage the development of reward cue attraction and pursuit (Anselme et al., 2013; Robinson et al., 2014, 2023) and habits (Dickinson, 1985). In these cases, behavioral responses to reward stimuli are heightened and are persistent. Here, the prediction error-like signaling at outcome that occurred over omission days could reflect an uncertainty about reward in the animals’ internal model of the task during omission sessions. However, inconsistent with this notion was the adjustment that rats made in their behavioral response to the cue, an adjustment that accorded with the new omission contingency. Further, in a prior study, animals with reward delivery experiences yoked exactly to animals undergoing the omission protocol did not exhibit similar behavioral learning consequences (Chang & Smith, 2011). Reward uncertainty therefore seems unlikely to explain our findings. A related factor, perceived scarcity of reward, might be a component of the positive error-like signal at reward occurrence, as scarcity can augment enjoyment (Johnson & Gallagher, 2011; Sehnert et al., 2014; Sevilla & Redden, 2014).

Reinforcement learning calculations, such as those in model-based and model-free accounts, do capture many aspects of sign-tracking and incentive salience (Dayan & Berridge, 2014), as well as its basis in phasic dopamine activity, including its rapid sensitivity to state changes and outcome value before the cue response becomes ingrained (Amaya et al., 2020; Chang et al., 2017; Dayan & Berridge, 2014; Derman et al., 2018; Robinson & Berridge, 2013; Tindell et al., 2009). Dopamine dynamics can also interface with model-based behavior (Gardner et al., 2018). As such, the role of dopamine in sign-tracking might initially be one rooted in model-based assessments for flexibility in reward value expectation and behavioral pursuit, but then transition to one rooted in model-free assessments to achieve a relatively habit-like inflexibility to changes of expected reward value. In principle, this shift would explain the sign-tracking behavior that animals exhibit after limited versus extended experience with the cue-reward relationship seen here (i.e., acquisition). However, it does not at face level explain the dopamine dynamics that occurred at the cue. Specifically, the entirely unchanged dopamine peak at cue occurrence across omission learning days seems inconsistent with model-based and model-free predictions, which ought to have ultimately affected cue-related dopamine release if it encoded reward prediction.

The dopamine dynamics we observed at the outcome time is more consistent with such learning models. To be consistent with these models, it would be as though at the time of the cue animals have a semi-accurate model of outcomes, whereas at the time of the outcome itself animals are continuously surprised by reward receipt or loss as though their predictions were always in error. A caveat to this possibility is the growth in either direction of the outcome related signals. Since these outcome-related dopamine signals if anything strengthened over sessions in our study here, and were unaffected on the trial level by outcomes received on the prior trail, this could be the model-free system’s attempt at competing with the robust strength of the model-based incentive salience signal at the cue onset. A purely learning account of the results might thus require a temporal separation of model-based (cue-related) and model-free (reward-related) calculations that compete for influence (Robke et al., 2024). This dual-system basis of learning has been accounted for in previous models (Lesaint et al., 2015, 2014) and may better represent the sign-tracking behavioral phenomenon than only either model-free or model-based learning. By using a combination of both systems, the model can encompass a larger swath of variable behaviors, as well as consider the internal states of the animals. What remains to be seen is how a model-based calculation at the cue can be influenced by model-free outcome signals such that cue-evoked behaviors change, yet the overall (often incorrect) prediction that reward will occur does not change. Arguably, a latent state-based account could succeed in explaining these results collectively. If the stable cue-evoked dopamine release reflects a stable internal model of the task during omission sessions, then it would imply that animals’ model of the task world involves a deep well of different cue-evoked behavior options that can be flexibly called forth when the cue arrives in a model-based manner.

The adjusted net contingency for causal relations (ANCCR) framework of associative learning (Jeong et al., 2022) is a potential candidate model of the dopamine transmission and the behavioral adjustments that occur as animals undergo omission. This assumes the animal can retrospect on a memory of the rewarding outcome (or perhaps, the lack of that outcome in omission) and infer the cause of it. It could be that the outcome signal growth that is observed across sessions is a representation of changes to the animals’ causal inference as they must consider their behavior and adjust to this new contingency to continue receiving these meaningful “causal targets”.

However, the above paragraphs suggest that a reinforcement learning account of our results is inadequate, or at least that it requires considerable shoehorning to fit. In contrast, we favor an incentive salience account. In this account, the reward cues acquire attractiveness with a neural representation in phasic dopamine activity. With time, the cue attractiveness becomes ingrained enough to decouple from the outcomes themselves, resulting in a persistence of motivation that is independent from learning and memory of rewards (Berridge, 2023). For its part, phasic dopamine itself seems to encode the persistent attractive features of cues while also encoding outcome updates when they occur. How dopamine at the time of reward cues becomes so misaligned from dopamine at the time of reward outcomes is a question of interest mechanistically, but it does appear to encode motivational persistence as seen here.

## Methods & Materials

### Subjects

16 PN 70-90 male (n = 8) and female (n = 8) Long Evans (Charles River) rats in conditioned reinforcement experiments and 9 PN 70-90 male (n = 3) and female (n = 6) Long Evans rats (Charles River) in fiber photometry experiments were single housed, and on a 12h light/dark cycle (lights on at 07:00). Rats were food restricted (7-15g of standard chow per day) to 85% of their free-feeding weight throughout testing. Water was available ad libitum. All procedures were approved by the Dartmouth College Institutional Animal Care and Use Committee.

### Testing apparatus

Tests were conducted in Med Associates chambers (20×30.5×29 cm) contained in sound- and light-attenuating cabinets (62×56×56 cm) equipped with a fan for airflow and background noise (∼68 dB). Chambers were illuminated by a house light on the back wall. Chambers contained two retractable levers on either side of a recessed magazine in which food rewards were delivered. For fiber photometry experiments, a chamber’s roof was removed and replaced with plexiglass walls that extended to the height of the chamber’s cabinet. A hole was drilled through the top of the cabinet for patch cable entry into the chamber. Lever deflections and magazine entries were recorded using the MED-PC IV software. Videos were recorded for behavioral analysis.

### Surgical procedures

Rats in the fiber photometry experiments were anesthetized with isoflurane gas and placed in a stereotaxic apparatus (Stoelting). All surgical procedures were conducted under aseptic conditions. A 5 µL, 33-gauge needle-tipped syringe (World Precision Instruments) was lowered to the target NAc site (from bregma: A/P +1.3; M/L ± 1.3; D/V -7.0; coordinates adapted from Flagel et al., 2011) and allowed to rest for 5 min. 0.8 µL of the viral vector containing the genetic material for GRAB_DA_ (Sun et al., 2020; 1×10^13^ vg/mL; pAAV-hsyn-GRAB_DA2m; Addgene) was delivered at a rate of 0.15 µL. Hemispheres were counterbalanced among rats. Following viral infusion, the needle was again allowed to rest for 5 min to allow viral dispersion. Optic fibers (400 µm diameter, numerical aperture (NA) = 0.5) were custom made and connected to a stainless-steel cannula (Thorlabs; model FP400URT). Fibers were implanted directly above the NAc viral infusion site and cemented to the skull using C&B Metabond (Parkell). Rats were allowed to recover in their cage for 3 weeks with daily injections of ketoprofen (dose mg/kg) and enrofloxacin (dose mg/kg) in 0.9% saline for the first 3 post-operative days.

### Sign-tracking acquisition training (Pavlovian conditioning)

Training began with a 30-minute magazine acclimation session where one pellet was delivered approximately every 30 seconds. Rats then received 12 days of Pavlovian sign-tracking (ST) acquisition training sessions. ST acquisition sessions contained 25 CS+ trials in which a 10-second presentation of a retractable lever was followed by 2, 45 mg grain pellet (BioServ) delivery, and 25 CS- trials in which the 10-second presentation of the other retractable lever was followed by nothing. CS+ and CS- lever sides were counterbalanced across animals. Trials were pseudorandomized so that no more than two of the same trial type were followed in sequence. Intertrial intervals were variable (ranging from 45 to 75 s) with an average length of 1 min.

Sessions lasted approximately 1 hour. Animals were tethered by their implant to a patch cable during the entirety of each session.

### Omission (negative automaintenance)

After completing 12 training days, one group of rats underwent 5 days of omission. Like the Pavlovian training schedule, these sessions contained 25, 10-second CS+ trials and 25, 10-second CS- trials. Under the omission condition, a deflection of the CS+ lever at any time during the trial would cancel reward delivery for that trial.

Pellets would be delivered only following CS+ trials in which the rats withheld lever deflections. Animals were tethered by their implant to a patch cable during the entirety of each session.

### Conditioned reinforcement

Training proceeded similarly as the sign-tracking acquisition procedures above, however, only one lever was presented (the reward-predictive, CS+ lever) and animals were not tethered. These sessions consisted of 25 CS+ trials, and lasted approximately 30 min. After the 12 sessions of training, one group of animals underwent 1 conditioned reinforcement session. In this session, a second lever on the opposite side of the magazine was presented to the rat for the duration of the session. When this lever was pressed, it would retract into the wall and the CS+ lever would be presented to the animal. No pellets were delivered during this session.

Another group of animals continued to 5 sessions of omission training with only 1, CS+ lever. Following these sessions, they underwent a conditioned reinforcement session as described previously. Animals were not tethered to a patch cable during these tasks.

### Fiber photometry recording

To assess dopamine responses to task events, GRAB_DA_ signals were recorded on the 1^st^ and final (12^th^) sessions of ST acquisition training, and all 5 sessions of omission. GRAB_DA_ fluorescence recording was completed using two wavelengths of light at 465 nm (excitation) and 405 nm (isosbestic) from two single wavelength LEDs that were driven by Doric Neuroscience Studio software. These two wavelengths were routed through a dichroic mirror (Doric) and into a lightweight metal jacketed, low autofluorescence patch cable (Doric, 400 µm, 0.57 NA) secured to the animal’s optic fiber implant using a ceramic mating sleeve (Doric). A dichroic mirror separated excitation and emission fluorescence carried from the fiber, and fluorescence was delivered to a photoreceiver (Newport model 2151, Doric). Fluorescent recordings were collected at 1017.2 Hz using Doric Neuroscience Studio software. Lock-in demodulation separated excitation and isosbestic fluorescence, and behavioral timestamps were recorded using TTL inputs.

### Fiber photometry preprocessing

Raw data from excitation and isosbestic channels were passed through a lowpass 5 Hz butterworth filter, and the signal was downsampled to 60 Hz. A least-squares linear fit was applied to the isosbestic to correct for photobleaching decay and align it to the excitation signal. ΔF/F was calculated by subtracting the fitted isosbestic signal from the excitation signal and dividing this over the fitted isosbestic signal. Z-scores were calculated by subtracting the mean ΔF/F across the entire session and dividing by the standard deviation to allow for comparisons across animals.

TTL timestamps for lever presentations and CS+ lever presses were used to creating analysis windows for relevant behavioral events. Minimum and maximum peak values were generated using the built-in min and max functions in Python. Area under the curve was calculated using a custom bidirectional trapezoidal sum function created in Python.

### Video analysis

Videos recorded on the first and final sign-tracking acquisition sessions and omission sessions were analyzed using DeepLabCut (version 2.1.10.4; Mathis et al., 2018). Each raw video was processed using Adobe Premiere Pro to reduce noise and enhance contrast for improved video quality. Approximately 200 frames were extracted for training. Labels were created for the snout, right and left ears, left and right levers, and the magazine. Frames with a likelihood higher than or equal to 0.80 were included in analysis. Location heatmaps only account for frames in which the CS+ lever was available to showcase the average location of a rat during CS+ presentations. To reduce sampling noise and produce a visually interpretable density, we applied a Gaussian smooth using a separable kernel (1D Gaussian convolved along rows and columns per bin) to yield a continuous-looking occupancy field without artificially broadening features excessively.

### Statistical analysis

The timing and number of lever deflections, magazine entries, and amount of time spent in the magazine area were recorded through Med-PC (Med Associates, St Albans, VT) and processed in Python. All statistical tests and linear regressions on both behavioral and fiber photometry recording data were completed in Python 3 (ttest_ind and ttest_rel functions from scipy-stats; LinearRegression function from sklearn), except for linear mixed effects (LME) models, which were run in R (packages: lme4; lmerTest). For all LME models, parameter estimates (est; β values), 95% confidence intervals (CI), and p-values are reported for the predictors. LMEs were chosen as they consider aspects of the data structure that repeated measures ANOVA cannot and allows for safer generalization to larger populations. All plots were created in Python (packages: matplotlib; seaborn) and designed in Adobe Illustrator.

## Acknowledgements

We thank Dr. Mitchell Spring for fiber photometry preprocessing assistance, Dr. Marvin Maechler for statistical and analysis consultation, as well as Drs. Katherine Nautiyal, Matthijs van der Meer, Talia Lerner, and Kate Wassum for their helpful comments and suggestions. We also thank Angela Shang for all figure illustrations.

## Data availability

The data in this study are available from the corresponding author upon request.

## Code availability

Fiber photometry preprocessing code is available at https://github.com/mspring-dartmouth/DoricProcessingTools. All other code developed in this study is available upon request.

## Author contributions

E.S.T. and K.S.S. developed the project and designed the experiments. E.S.T. and D.G. performed the experiments and analyzed the data. E.S.T. wrote the manuscript, with contribution and edits from K.S.S. and D.G.

## References

1. Ahrens, A. M., Meyer, P. J., Ferguson, L. M., Robinson, T. E., & Aldridge, J. W. (2016). Neural Activity in the Ventral Pallidum Encodes Variation in the Incentive Value of a Reward Cue. The Journal of Neuroscience: The Official Journal of the Society for Neuroscience, 36(30), 7957–7970.

2. Amaya, K. A., Stott, J. J., & Smith, K. S. (2020). Sign-tracking behavior is sensitive to outcome devaluation in a devaluation context-dependent manner: implications for analyzing habitual behavior. Learning & Memory , 27(4), 136–149.

3. Anselme, P., Robinson, M. J. F., & Berridge, K. C. (2013). Reward uncertainty enhances incentive salience attribution as sign-tracking. Behavioural Brain Research, 238, 53–61.

4. Berke, J. D. (2018). What does dopamine mean? Nature Neuroscience, 21(6), 787–793.

5. Berridge, K. C. (2004). Motivation concepts in behavioral neuroscience. Physiology & Behavior, 81(2), 179–209.

6. Berridge, K. C. (2007). The debate over dopamine’s role in reward: the case for incentive salience. Psychopharmacology, 191(3), 391–431.

7. Berridge, K. C. (2023). Separating desire from prediction of outcome value. Trends in Cognitive Sciences, 27(10), 932–946.

8. Breland, K., & Breland, M. (1961). The misbehavior of organisms. The American Psychologist, 16(11), 681–684.

9. Brown, P. L., & Jenkins, H. M. (1968). Auto-shaping of the pigeon’s key-peck. Journal of the Experimental Analysis of Behavior, 11(1), 1–8.

10. Chang, S. E., Smedley, E. B., Stansfield, K. J., Stott, J. J., & Smith, K. S. (2017). Optogenetic Inhibition of Ventral Pallidum Neurons Impairs Context-Driven Salt Seeking. The Journal of Neuroscience: The Official Journal of the Society for Neuroscience, 37(23), 5670–5680.

11. Chang, S. E., & Smith, K. S. (2016). An omission procedure reorganizes the microstructure of sign-tracking while preserving incentive salience. Learning & Memory , 23(4), 151–155.

12. Costa, K. M., & Schoenbaum, G. (2022). Dopamine. Current Biology: CB, 32(15), R817–R824.

13. Davey, G. C., Oakley, D., & Cleland, G. G. (1981). Autoshaping in the rat: Effects of omission on the form of the response. Journal of the Experimental Analysis of Behavior, 36(1), 75–91.

14. Daw, N. D., Niv, Y., & Dayan, P. (2005). Uncertainty-based competition between prefrontal and dorsolateral striatal systems for behavioral control. Nature Neuroscience, 8(12), 1704–1711.

15. Day, J. J., Roitman, M. F., Wightman, R. M., & Carelli, R. M. (2007). Associative learning mediates dynamic shifts in dopamine signaling in the nucleus accumbens. Nature Neuroscience, 10(8), 1020–1028.

16. Day, J. J., Wheeler, R. A., Roitman, M. F., & Carelli, R. M. (2006). Nucleus accumbens neurons encode Pavlovian approach behaviors: evidence from an autoshaping paradigm. The European Journal of Neuroscience, 23(5), 1341–1351.

17. Dayan, P., & Berridge, K. C. (2014). Model-based and model-free Pavlovian reward learning: revaluation, revision, and revelation. Cognitive, Affective & Behavioral Neuroscience, 14(2), 473–492.

18. Derman, R. C., Schneider, K., Juarez, S., & Delamater, A. R. (2018). Sign-tracking is an expectancy-mediated behavior that relies on prediction error mechanisms. Learning & Memory , 25(10), 550–563.

19. Dickinson, A. (1985). Actions and habits: the development of behavioural autonomy. Philosophical Transactions of the Royal Society of London, 308(1135), 67–78.

20. Edwards, S., & Koob, G. F. (2010). Neurobiology of dysregulated motivational systems in drug addiction. Future Neurology, 5(3), 393–401.

21. Enriquez-Traba, J., Arenivar, M., Yarur-Castillo, H. E., Noh, C., Flores, R. J., Weil, T., Roy, S., Usdin, T. B., LaGamma, C. T., Wang, H., Tsai, V. S., Kerspern, D., Moritz, A. E., Sibley, D. R., Lutas, A., Moratalla, R., Freyberg, Z., & Tejeda, H. A. (2024). Dissociable control of motivation and reinforcement by distinct ventral striatal dopamine receptors. Nature Neuroscience, 28(1), 105–121.

22. Ferguson, L. M., Ahrens, A. M., Longyear, L. G., & Aldridge, J. W. (2020). Neurons of the ventral tegmental area encode individual differences in motivational “wanting” for reward cues. The Journal of Neuroscience: The Official Journal of the Society for Neuroscience, 40(46), 8951–8963.

23. Flagel, S. B., Akil, H., & Robinson, T. E. (2009). Individual differences in the attribution of incentive salience to reward-related cues: Implications for addiction. Neuropharmacology, 56 *Suppl 1*, 139–148.

24. Flagel, S. B., Clark, J. J., Robinson, T. E., Mayo, L., Czuj, A., Willuhn, I., Akers, C. A., Clinton, S. M., Phillips, P. E. M., & Akil, H. (2011). A selective role for dopamine in stimulus-reward learning. Nature, 469(7328), 53–57.

25. Flagel, S. B., & Robinson, T. E. (2017). Neurobiological Basis of Individual Variation in Stimulus-Reward Learning. Current Opinion in Behavioral Sciences, 13, 178–185.

26. Fraser, K. M., Collins, V., Wolff, A. R., Ottenheimer, D. J., Bornhoft, K. N., Pat, F., Chen, B. J., Janak, P. H., & Saunders, B. T. (2025). Contextual cues facilitate dynamic value encoding in the mesolimbic dopamine system. Current Biology: CB, 35(4), 746–760.e5.

27. Fraser, K. M., & Janak, P. H. (2017). Long-lasting contribution of dopamine in the nucleus accumbens core, but not dorsal lateral striatum, to sign-tracking. The European Journal of Neuroscience, 46(4), 2047–2055.

28. Gardner, M. P. H., Schoenbaum, G., & Gershman, S. J. (2018). Rethinking dopamine as generalized prediction error. *Proceedings*. Biological Sciences, 285(1891), 20181645.

29. Garr, E., Cheng, Y., Jeong, H., Brooke, S., Castell, L., Bal, A., Magnard, R., Namboodiri, V. M. K., & Janak, P. H. (2023). Mesostriatal dopamine is sensitive to specific cue-reward contingencies. In bioRxiv (p. 2023.06.05.543690). 10.1101/2023.06.05.543690

30. Goedhoop, J., Arbab, T., & Willuhn, I. (2023). Anticipation of Appetitive Operant Action Induces Sustained Dopamine Release in the Nucleus Accumbens. The Journal of Neuroscience: The Official Journal of the Society for Neuroscience, 43(21), 3922–3932.

31. Hamid, A. A., Pettibone, J. R., Mabrouk, O. S., Hetrick, V. L., Schmidt, R., Vander Weele, C. M., Kennedy, R. T., Aragona, B. J., & Berke, J. D. (2016). Mesolimbic dopamine signals the value of work. Nature Neuroscience, 19(1), 117–126.

32. Holland, P. C., Asem, J. S. A., Galvin, C. P., Keeney, C. H., Hsu, M., Miller, A., & Zhou, V. (2014). Blocking in autoshaped lever-pressing procedures with rats. Learning & Behavior, 42(1), 1–21.

33. Howe, M. W., Tierney, P. L., Sandberg, S. G., Phillips, P. E. M., & Graybiel, A. M. (2013). Prolonged dopamine signalling in striatum signals proximity and value of distant rewards. Nature, 500(7464), 575–579.

34. Iglesias, A. G., Chiu, A. S., Wong, J., Campus, P., Li, F., Liu, Z. N., Bhatti, J. K., Patel,

35. S. A., Deisseroth, K., Akil, H., Burgess, C. R., & Flagel, S. B. (2023). Inhibition of Dopamine Neurons Prevents Incentive Value Encoding of a Reward Cue: With Revelations from Deep Phenotyping. The Journal of Neuroscience: The Official Journal of the Society for Neuroscience, 43(44), 7376–7392.

36. Jeong, H., Taylor, A., Floeder, J. R., Lohmann, M., Mihalas, S., Wu, B., Zhou, M., Burke, D. A., & Namboodiri, V. M. K. (2022). Mesolimbic dopamine release conveys causal associations. *Science (New York*, N.Y.), 378(6626), eabq6740.

37. Johnson, A. W., & Gallagher, M. (2011). Greater effort boosts the affective taste properties of food. *Proceedings*. Biological Sciences, 278(1711), 1450–1456.

38. Keefer, S. E., Kochli, D. E., & Calu, D. J. (2022). Inactivation of the basolateral amygdala to insular cortex pathway makes sign-tracking sensitive to outcome devaluation. ENeuro, 9(5), ENEURO.0156-22.2022.

39. Kochli, D. E., Keefer, S. E., Gyawali, U., & Calu, D. J. (2020). Basolateral amygdala to nucleus accumbens communication differentially mediates devaluation sensitivity of sign- and goal-tracking rats. Frontiers in Behavioral Neuroscience, 14, 593645.

40. Lerner, T. N., Holloway, A. L., & Seiler, J. L. (2021). Dopamine, Updated: Reward Prediction Error and Beyond. Current Opinion in Neurobiology, 67, 123–130.

41. Lesaint, F., Sigaud, O., Clark, J. J., Flagel, S. B., & Khamassi, M. (2015). Experimental predictions drawn from a computational model of sign-trackers and goal-trackers. Journal of Physiology, Paris, 109(1–3), 78–86.

42. Lesaint, F., Sigaud, O., Flagel, S. B., Robinson, T. E., & Khamassi, M. (2014). Modelling individual differences in the form of Pavlovian conditioned approach responses: a dual learning systems approach with factored representations. PLoS Computational Biology, 10(2), e1003466.

43. Locurto, C., Terrace, H. S., & Gibbon, J. (1976). Autoshaping, random control, and omission training in the rat. Journal of the Experimental Analysis of Behavior, 26(3), 451–462.

44. María-Ríos, C. E., Fitzpatrick, C. J., Czesak, F. N., & Morrow, J. D. (2023). Effects of predictive and incentive value manipulation on sign- and goal-tracking behavior. Neurobiology of Learning and Memory, 203, 107796.

45. Mathis, A., Mamidanna, P., Cury, K. M., Abe, T., Murthy, V. N., Mathis, M. W., & Bethge, M. (2018). DeepLabCut: markerless pose estimation of user-defined body parts with deep learning. Nature Neuroscience, 21(9), 1281–1289.

46. McClure, S. M., Daw, N. D., & Montague, P. R. (2003). A computational substrate for incentive salience. Trends in Neurosciences, 26(8), 423–428.

47. Mohebi, A., Pettibone, J. R., Hamid, A. A., Wong, J.-M. T., Vinson, L. T., Patriarchi, T., Tian, L., Kennedy, R. T., & Berke, J. D. (2019). Dissociable dopamine dynamics for learning and motivation. Nature, 570(7759), 65–70.

48. Niv, Y., Daw, N. D., Joel, D., & Dayan, P. (2007). Tonic dopamine: opportunity costs and the control of response vigor. Psychopharmacology, 191(3), 507–520.

49. Robinson, M. J. F., Anselme, P., Fischer, A. M., & Berridge, K. C. (2014). Initial uncertainty in Pavlovian reward prediction persistently elevates incentive salience and extends sign-tracking to normally unattractive cues. Behavioural Brain Research, 266, 119–130.

50. Robinson, M. J. F., & Berridge, K. C. (2013). Instant transformation of learned repulsion into motivational “wanting.” Current Biology: CB, 23(4), 282–289.

51. Robinson, M. J. F., Bonmariage, Q. S. A., & Samaha, A.-N. (2023). Unpredictable, intermittent access to sucrose or water promotes increased reward pursuit in rats. Behavioural Brain Research, 453(114612), 114612.

52. Robinson, T. E., & Berridge, K. C. (1993). The neural basis of drug craving: an incentive-sensitization theory of addiction. Brain Research. Brain Research Reviews, 18(3), 247–291.

53. Robinson, T. E., & Berridge, K. C. (2008). Review. The incentive sensitization theory of addiction: some current issues. *Philosophical Transactions of the Royal Society of London. Series B*, Biological Sciences, 363(1507), 3137–3146.

54. Robinson, T. E., & Berridge, K. C. (2025). The incentive-sensitization theory of addiction 30 years on. Annual Review of Psychology, 76(1), 29–58.

55. Robke, R., Arbab, T., Smith, R., & Willuhn, I. (2024). Value-driven adaptations of mesolimbic dopamine release are governed by both model-based and model-free mechanisms. ENeuro, 11(7), ENEURO.0223-24.2024.

56. Salamone, J. D., Correa, M., Farrar, A., & Mingote, S. M. (2007). Effort-related functions of nucleus accumbens dopamine and associated forebrain circuits. Psychopharmacology, 191(3), 461–482.

57. Saunders, B. T., & Robinson, T. E. (2012). The role of dopamine in the accumbens core in the expression of Pavlovian-conditioned responses. The European Journal of Neuroscience, 36(4), 2521–2532.

58. Saunders, B. T., & Robinson, T. E. (2013). Individual variation in resisting temptation: implications for addiction. Neuroscience and Biobehavioral Reviews, *37*(9 Pt A), 1955–1975.

59. Schultz, W. (1998). Predictive reward signal of dopamine neurons. Journal of Neurophysiology, 80(1), 1–27.

60. Sehnert, S., Franks, B., Yap, A. J., & Higgins, E. T. (2014). Scarcity, engagement, and value. Motivation and Emotion, 38(6), 823–831.

61. Sevilla, J., & Redden, J. P. (2014). Limited availability reduces the rate of satiation. JMR, Journal of Marketing Research, 51(2), 205–217.

62. Smith, K. S., Berridge, K. C., & Aldridge, J. W. (2011). Disentangling pleasure from incentive salience and learning signals in brain reward circuitry. Proceedings of the National Academy of Sciences of the United States of America, 108(27), E255–64.

63. Smith, K. S., Virkud, A., Deisseroth, K., & Graybiel, A. M. (2012). Reversible online control of habitual behavior by optogenetic perturbation of medial prefrontal cortex. Proceedings of the National Academy of Sciences of the United States of America, 109(46), 18932–18937.

64. Stiers, M., & Silberberg, A. (1974). Lever-contact responses in rats: automaintenance with and without a negative response-reinforcer dependency. Journal of the Experimental Analysis of Behavior, 22(3), 497–506.

65. Sun, F., Zhou, J., Dai, B., Qian, T., Zeng, J., Li, X., Zhuo, Y., Zhang, Y., Wang, Y., Qian, C., Tan, K., Feng, J., Dong, H., Lin, D., Cui, G., & Li, Y. (2020). Next-generation GRAB sensors for monitoring dopaminergic activity in vivo. Nature Methods, 17(11), 1156–1166.

66. Tindell, A. J., Smith, K. S., Berridge, K. C., & Aldridge, J. W. (2009). Dynamic computation of incentive salience: “wanting” what was never “liked.” The Journal of Neuroscience: The Official Journal of the Society for Neuroscience, 29(39), 12220–12228.

67. Townsend, E. S., Amaya, K. A., Smedley, E. B., & Smith, K. S. (2023). Nucleus accumbens core acetylcholine receptors modulate the balance of flexible and inflexible cue-directed motivation. Scientific Reports, 13(1), 13375.

68. Townsend, E. S., & Smith, K. S. (2025). Behavioral microanalyses refine sign-tracking characterization and uncover different response dynamics during omission and extinction learning. Learning & Memory, 32(3), a054065.

69. Tunstall, B. J., & Kearns, D. N. (2015). Sign-tracking predicts increased choice of cocaine over food in rats. Behavioural Brain Research, 281, 222–228.

70. Warlow, S. M., Naffziger, E. E., & Berridge, K. C. (2020). The central amygdala recruits mesocorticolimbic circuitry for pursuit of reward or pain. Nature Communications, 11(1), 2716.

71. Williams, D. R., & Williams, H. (1969). Auto-maintenance in the pigeon: sustained pecking despite contingent non-reinforcement. Journal of the Experimental Analysis of Behavior, 12(4), 511–520.

72. Wyvell, C. L., & Berridge, K. C. (2000). Intra-accumbens amphetamine increases the conditioned incentive salience of sucrose reward: enhancement of reward “wanting” without enhanced “liking” or response reinforcement. The Journal of Neuroscience: The Official Journal of the Society for Neuroscience, 20(21), 8122–8130.

73. Zhang, J., Berridge, K. C., Tindell, A. J., Smith, K. S., & Aldridge, J. W. (2009). A neural computational model of incentive salience. PLoS Computational Biology, 5(7), e1000437.

