## Supplemental Figures for "Phasic dopamine encodes persistent attraction to reward cues"

### Supplemental Material

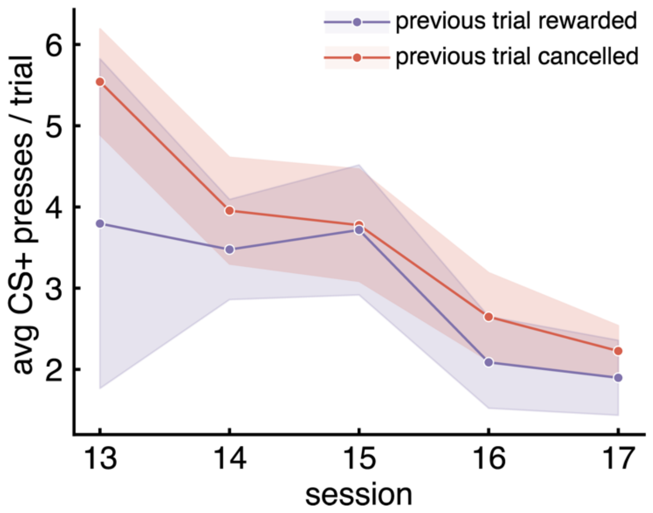


**Supplemental Figure 1: Reward receipt or cancellation does not affect sign-tracking on the subsequent trial.** Average CS+ lever presses on a given trial during omission (sessions 13-17) when the previous trial reward was delivered (red) or cancelled (purple). No effect of previous trial type (reward delivered or cancelled) was seen, but a significant effect of session was found (est: -0.73; CI: -1.01 – -0.44; p<0.001). Error ribbons represent ± SEM.

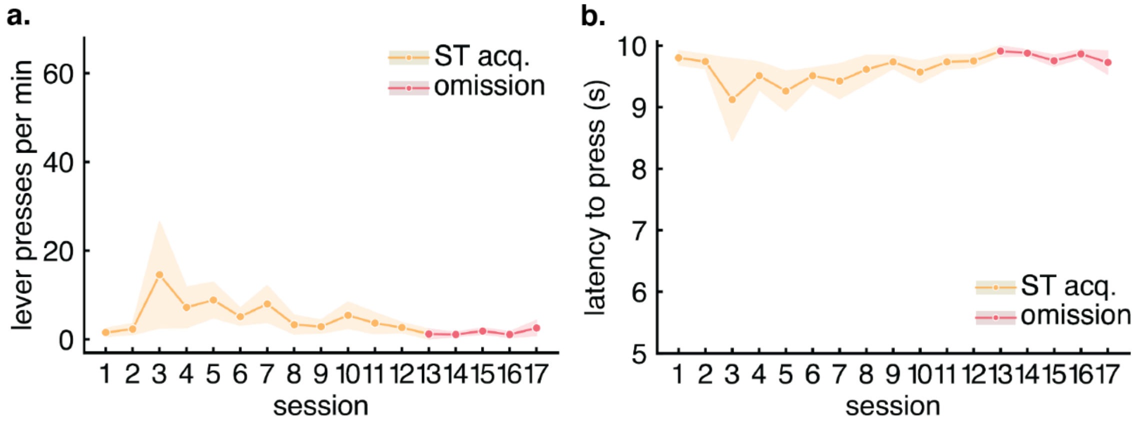


**Supplemental Figure 2: CS- lever-directed behaviors during sign-tracking acquisition and omission. A)** CS- lever presses per minute during sign-tracking acquisition training (sessions 1-12; teal) and omission (sessions 13-17, pink). **B)** Latency to press the CS- in seconds, when it is pressed, during sign-tracking acquisition training (sessions 1-12; teal) and omission (sessions 13-17, pink). For all plots, error ribbons represent ± SEM.

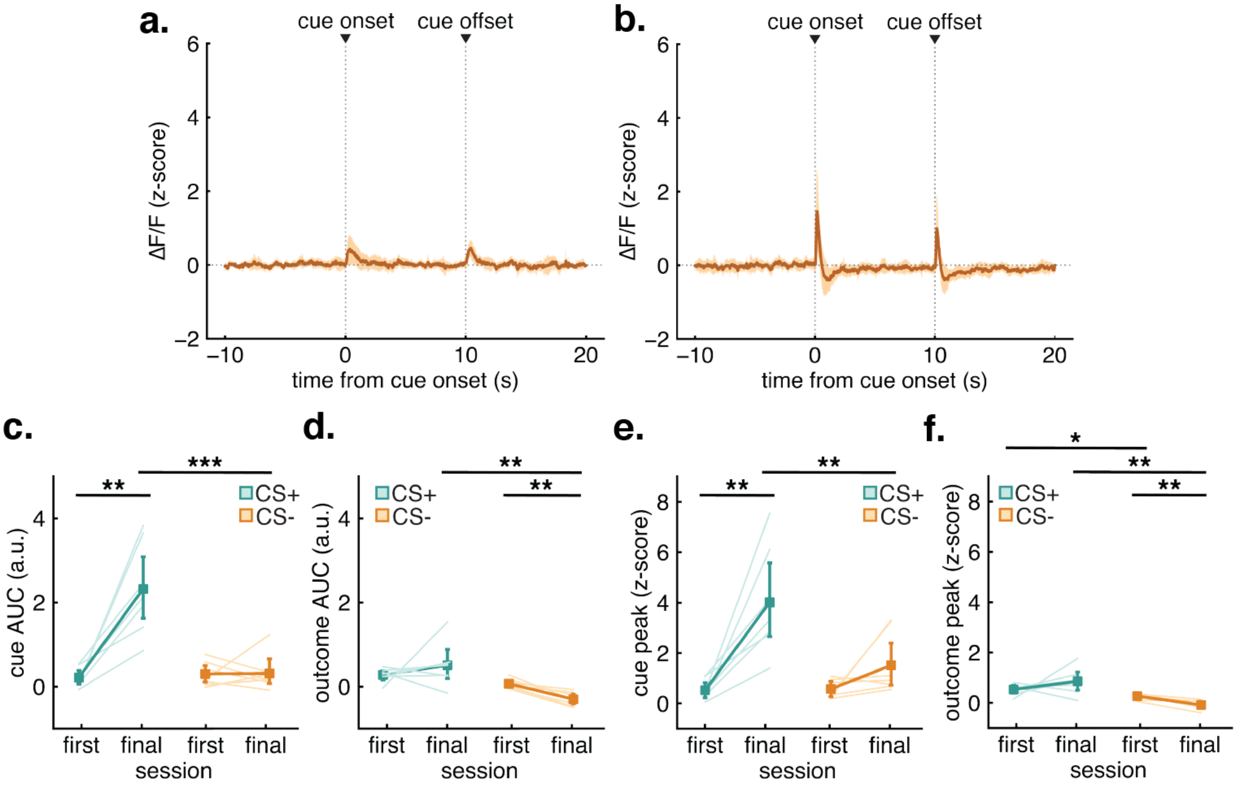


**Supplemental Figure 3: DA dynamics during CS- trials. A)** Average ΔF/F of the 25 CS- trials in the first sign-tracking acquisition session for the 10 s prior, 10 s during, and 10 s following the CS- onset (0 s) and offset (10 s) are denoted by dotted lines. **B)** Average ΔF/F of the 25 CS- trials in the final sign-tracking acquisition session. **C)** AUC (arbitrary units; a.u.) of DA activity comparing CS+ (teal) and CS- (orange) presentations. CS+ evoked signaling was significantly greater than that of CS- evoked signaling during the final acquisition session ($t=5.41$; $p=0.001$). **D)** AUC of DA during the outcome period following the CS+ (teal; reward occurs) and CS- (orange; no reward occurs). Outcome-evoked signaling was significantly greater following the CS+ rather than the CS- during the final acquisition session ($t=3.63$; $p=0.01$). CS- outcome-evoked signaling also dropped significantly from the first to final sessions ($t=-5.94$; $p=0.002$). **E)** Peak z-score during the 1 s time bin beginning at the CS+ (teal) and CS- (orange) onset. CS+ evoked peaks were significantly greater than those of CS- evoked signaling during the final acquisition session ($t=5.95$; $p<0.001$). **F)** Peak z-score during the outcome period. Outcome-evoked peaks were significantly greater following the CS+ rather than the CS- during both the first ($t=2.87$; $p=0.029$) and final acquisition session ($t=3.88$; $p=0.008$). Outcome signal peaks following the CS- dropped significantly from the first to final acquisition sessions ($t=-4.67$; $p=0.003$). For all plots, error ribbons and bars represent ± SEM. Asterisks (*) represent significant results.
